# A Fractal-Dimension Framework for Quantifying Self-Similarity in Chromatin Folding

**DOI:** 10.64898/2026.05.06.723123

**Authors:** Anass El-Yaagoubi, Ali Balubaid, Moo Chung, Jesper Tegnér, Hernando Ombao

## Abstract

The three-dimensional folding of DNA is essential for genome function, but its organization remains difficult to summarize quantitatively across genomic scales. Here, we study DNA folding from Hi-C contact data using a network-based notion of fractal dimension. In this representation, genomic loci are treated as nodes, and observed Hi-C contacts define weighted edges, so that frequently interacting loci are closer in the resulting network. We then estimate fractal dimension using two complementary graph-based methods: the correlation dimension and the sandbox dimension.

Validation on synthetic networks shows that the proposed estimators detect clear scaling behavior in hierarchical fractal-like networks, while distinguishing them from networks with local clustering but no stable multiscale self-similarity. Applied to intrachromosomal Hi-C data from the IMR90 human cell line, the method reveals approximate linear scaling regimes on log–log plots, suggesting fractal-like organization in chromatin contact networks. At the chromosome level, estimated fractal dimension tends to increase with chromosome size: larger chromosomes often have dimensions closer to 3, consistent with more compact and space-filling organization, whereas shorter chromosomes tend to have lower dimensions, closer to 1, consistent with simpler and more open folding patterns. A sliding-window analysis at 5 kb resolution further shows that fractal organization varies substantially along chromosomes rather than remaining uniform across genomic position.

These results suggest that graph-based fractal dimension provides an interpretable summary of DNA folding complexity at both global and local scales. More broadly, the proposed framework offers a quantitative way to study multiscale genome organization from Hi-C data using tools from network geometry.

## 1 Introduction

Every living organism carries a genome that governs development, regulates physiological activity, supports reproduction, and ultimately shapes survival and lifespan [1]. This genome is encoded in deoxyribonucleic acid (DNA), the molecule that stores the genetic information required for cellular function [2]. Alterations in the DNA sequence can substantially increase the risk of disease [3, 4]. At the same time, biological activity is not determined by sequence alone. The three-dimensional conformation of DNA also plays a fundamental role in regulating gene expression and other essential cellular processes [5, 6].

Characterizing DNA organization is challenging because of the extraordinary disparity in scale between genome length and nuclear volume, as illustrated in Figure 1. In human cells, the genome contains roughly 3 billion nucleotides and, if fully extended, would measure on the order of two meters, yet it is compacted within a nucleus whose diameter is only a few micrometers. This remarkable level of packaging is achieved through chromatin folding, whereby DNA and associated proteins are arranged into a highly compact and dynamic structure [7]. Chromatin folding is not merely a mechanism for storage. It is also closely tied to genome regulation, since the formation of loops, domains, and higher-order structures influences the accessibility and coordination of genomic elements [8, 9]. Understanding the principles that govern this folding remains a major challenge because chromatin architecture is complex, multiscale, and spatially heterogeneous.

**Figure 1:**
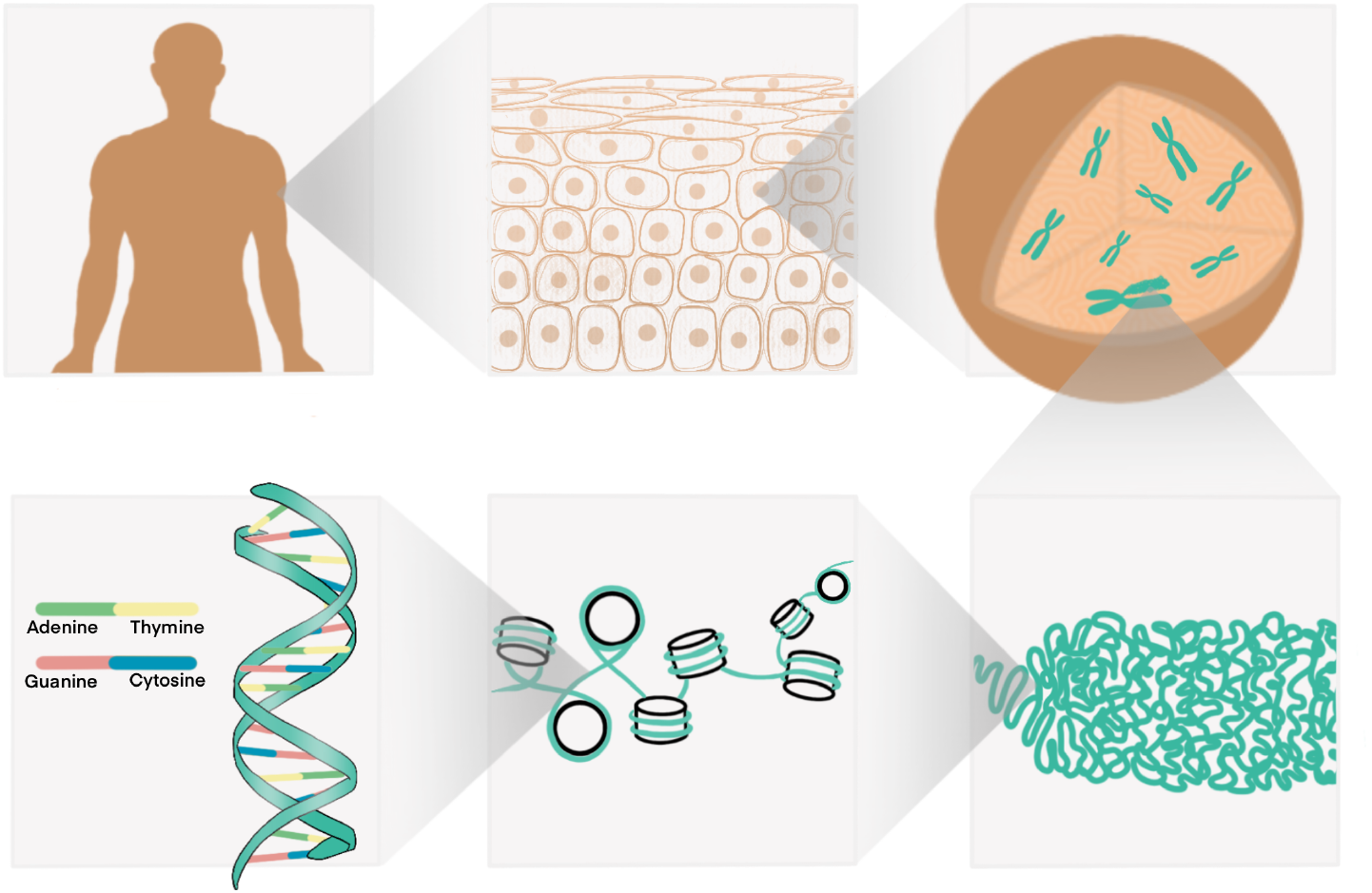
Various scales of biological organization, from the human body to the cell, nucleus, folded chromosomes, local chromatin domains, and the DNA sequence.

Recent advances in sequencing technologies have greatly expanded our ability to study genome organization in three dimensions [10, 11]. In particular, chromosome conformation capture methods measure how often different regions of the genome are found close to one another inside the nucleus. Hi-C is a genome-wide version of this approach: DNA is first fixed in its three-dimensional state, nearby fragments are ligated together, and the resulting paired fragments are sequenced to identify which genomic loci were spatially proximal [12]. The output is a contact map, where each entry records the frequency of interaction between two genomic loci. Larger contact frequencies are generally interpreted as evidence that the corresponding loci are more often close in three-dimensional space. These contact maps provide genome-wide information on chromatin architecture and have revealed important structural features, including compartments, topologically associating domains (TADs), and chromatin loops [13]. Although highly informative, Hi-C data are also challenging to interpret because they reflect complex interaction patterns across many genomic scales and may contain missing or low-coverage regions, particularly in difficult-to-map genomic segments.

One natural question is whether chromatin folding exhibits approximate self-similar organization across scales. Self-similar patterns, often associated with fractal geometry, arise in many natural and biological systems, including river networks, coastlines, trees, lungs, vascular systems, and neuronal branching. Such structures often emerge from recursive growth, repeated local interactions, or simple organizing rules that generate complex large-scale behavior, and they are frequently associated with efficient space filling, transport, or exchange. This analogy has motivated the idea that chromatin may also possess fractal-like structural organization. If such multiscale regularity is present, then fractal dimension provides a natural quantitative tool for summarizing it. In the present work, we investigate whether chromatin interaction networks derived from Hi-C data exhibit scaling behavior consistent with fractal organization, and whether fractal dimension can serve as a quantitative summary of the underlying DNA folding complexity.

To investigate this question, we represent each chromosome as a weighted interaction network, where genomic loci are nodes and Hi-C contact frequencies define interaction-derived distances between them. This network view makes it possible to study chromatin folding using tools designed for complex multiscale systems. In particular, fractal dimension provides a compact way to summarize how interaction neighborhoods grow with scale, and can therefore be used to compare chromosomes, assess local variation along the genome, and distinguish more compact from more diffuse folding patterns. In this work, we focus on two graph-based estimators of fractal dimension: the correlation dimension and the sandbox dimension.

The main objective of this study is to develop and evaluate a network-based framework for estimating fractal dimension from Hi-C chromatin contact data. We first validate the approach on synthetic networks with contrasting structural organization, including networks with and without clear multiscale self-similarity. We then assess the stability and computational cost of the estimators, before applying the framework to human IMR90 intrachromosomal Hi-C data. This application examines chromosome-level scaling behavior, the relationship between chromosome size and estimated fractal dimension, and local heterogeneity through high-resolution sliding-window analysis. The remainder of the paper is organized as follows. Section 2 introduces the Hi-C setting, the chromatin network construction, and the fractal-dimension estimators. Section 3 presents the validation, uncertainty, and computational analyses. Section 4 reports the chromosome-level and local genomic results.

## 2 Methodology

This section describes the construction of chromatin interaction networks from Hi-C data and the graph-based fractal-dimension methods used to analyze them. We first summarize the Hi-C data structure, then define the weighted network representation of each chromosome, and finally introduce the correlation and sandbox estimators used in the empirical analysis.

### 2.1 Chromatin Networks

Chromatin conformation capture (3C) techniques are designed to characterize the three-dimensional organization of DNA in the nucleus. This is a challenging task because the genome is extremely long while the spatial scale of folding is very small. Hi-C, a genome-wide high-throughput chromosome conformation capture technique, addresses this difficulty by converting spatial DNA interactions into a contact matrix, where each entry records how often two loci are observed together in sequencing data. The main steps of the protocol are illustrated in Figure 2. In brief, chromatin is first crosslinked to preserve spatial proximity, then digested into fragments, labeled and ligated to capture nearby interactions, purified and amplified, and finally sequenced, yielding a genome-wide map of pairwise interaction frequencies.

**Figure 2:**
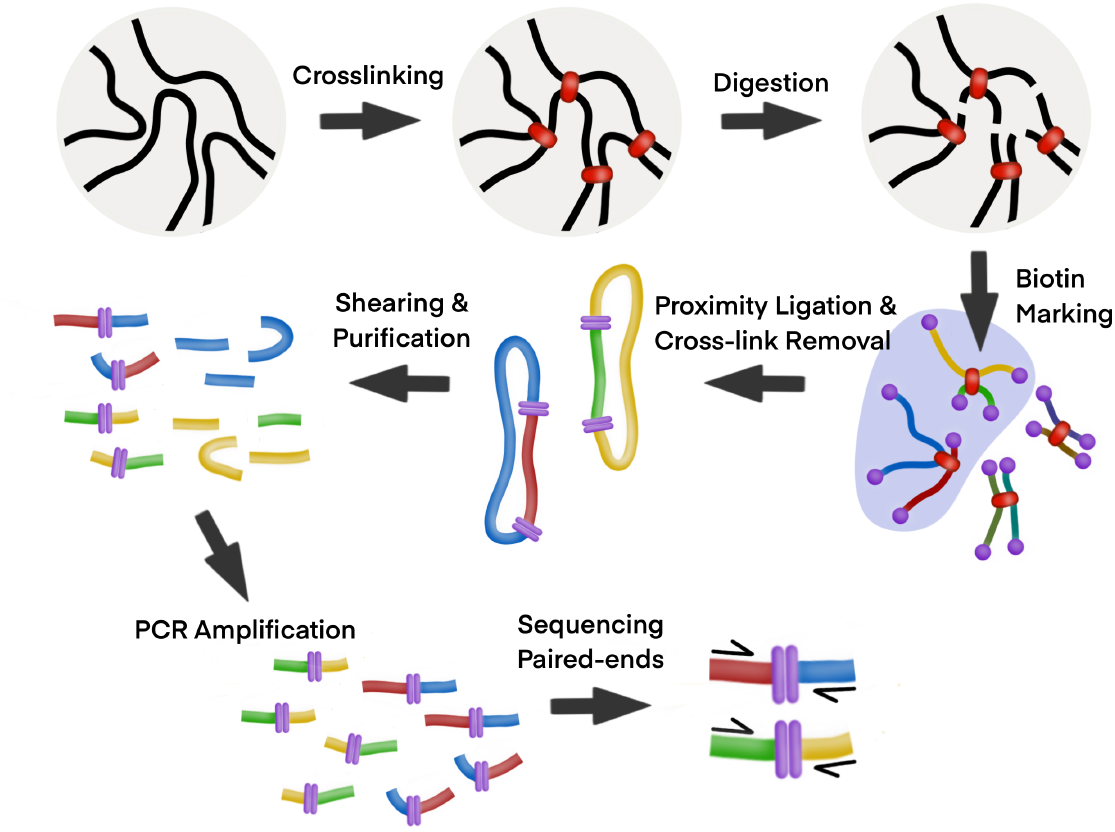
Illustration of the Hi-C workflow. The protocol involves crosslinking, digestion, biotin labeling, proximity ligation and reverse crosslinking, shearing and purification, PCR amplification, and high-throughput paired-end sequencing, resulting in a contact map of DNA loci interactions.

In this work, we analyze intrachromosomal Hi-C contact matrices from the IMR90 cell line obtained from the Gene Expression Omnibus (GEO) database of the National Center for Biotechnology Information (NCBI), accession GSE63525, associated with the high-resolution in situ Hi-C study of Rao et al. [14]. The data are organized in a per-chromosome folder structure and are available at multiple resolutions. We focus on 50 kb resolution for global chromosome-level fractal-dimension estimation, since this scale provide a useful balance between structural detail and computational feasibility. We also examine the 5 kb data, but only through a sliding-window analysis, as full chromosome-wide estimation at this resolution is substantially more demanding in terms of memory and computation. Throughout the analysis, we preferentially use the MAPQGE30 quality-filtered data. For the contact matrix itself, our working choice is the RAW observed track, while alternative matrix normalizations such as KR, VC, and SQRTVC are also available in the data source.

To represent chromatin organization as a graph, each chromosome is partitioned into contiguous genomic bins at a fixed resolution. The resulting chromatin network for chromosome *k* is written as

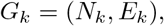

where *N*_*k*_ denotes the set of genomic loci and *E*_*k*_ denotes the set of weighted edges induced by observed Hi-C contacts. If 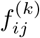 denotes the observed contact frequency between loci *i* and *j*, then an edge is introduced only when 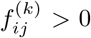. In particular, 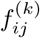 is treated as the absence of an edge. For pairs with positive contact frequency, we define the edge weight by

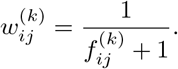

Under this construction, larger contact frequencies correspond to smaller edge weights and therefore to greater effective proximity in the graph.

The number of nodes in the network depends directly on chromosome length and resolution. Here, *kb* denotes a kilobase (1,000 base pairs). As the resolution becomes finer, the number of genomic bins increases substantially, which in turn increases the size of the graph and the computational cost of distance-based estimation procedures. For example, the largest human chromosome has length about 249 Mbp, whereas one of the smallest, chromosome 21, has length about 47 Mbp. This corresponds roughly to about 5,000 versus 940 bins at 50 kb resolution, about 10,000 versus 1,880 bins at 25 kb resolution, and about 50,000 versus 9,400 bins at 5 kb resolution. Since the Hi-C contact map is a square matrix, its size grows quadratically with the number of bins. If stored densely using 32-bit floating-point values, this corresponds approximately to 100 MB versus 3.5 MB at 50 kb, 400 MB versus 14 MB at 25 kb, and 10 GB versus 350 MB at 5 kb for a single chromosome. Thus, while 50 kb and 25 kb remain feasible for chromosome-level analysis on a standard laptop, full chromosome-wide estimation at 5 kb becomes much more demanding in both memory and runtime. For this reason, 5 kb data are not handled through full-chromosome global estimation; instead, they are analyzed using a sliding-window strategy. When a given window does not admit reliable estimation, for example because the induced graph is disconnected or does not provide a usable distance structure, the corresponding output is recorded as `NA` and the gap is retained rather than artificially filled.

### 2.2 Self-Similarity and Fractal Dimensions

In order for DNA to fit within the confined volume of the nucleus, it must fold in a highly organized manner. Although the mechanisms governing chromatin architecture are complex, an important working hypothesis is that chromatin organization may display approximate self-similarity across scales. Self-similarity refers to the persistence of structural patterns under changes of scale. Such patterns are widespread in nature. Figure 3 presents two familiar examples, lungs and a fern leaf, both of which exhibit repeated branching motifs across scales. Many other natural systems display related behavior, including river networks, coastlines, blood vessels, neurons, snowflakes, broccoli, and trees. In many cases, these structures arise from recursive growth rules or repeated local interactions, where comparatively simple mechanisms generate complex large-scale organization. Fractal-like architectures are also often associated with efficient space filling, transport, exchange, or coverage, which helps explain why they occur so frequently in biological and physical systems.

**Figure 3:**
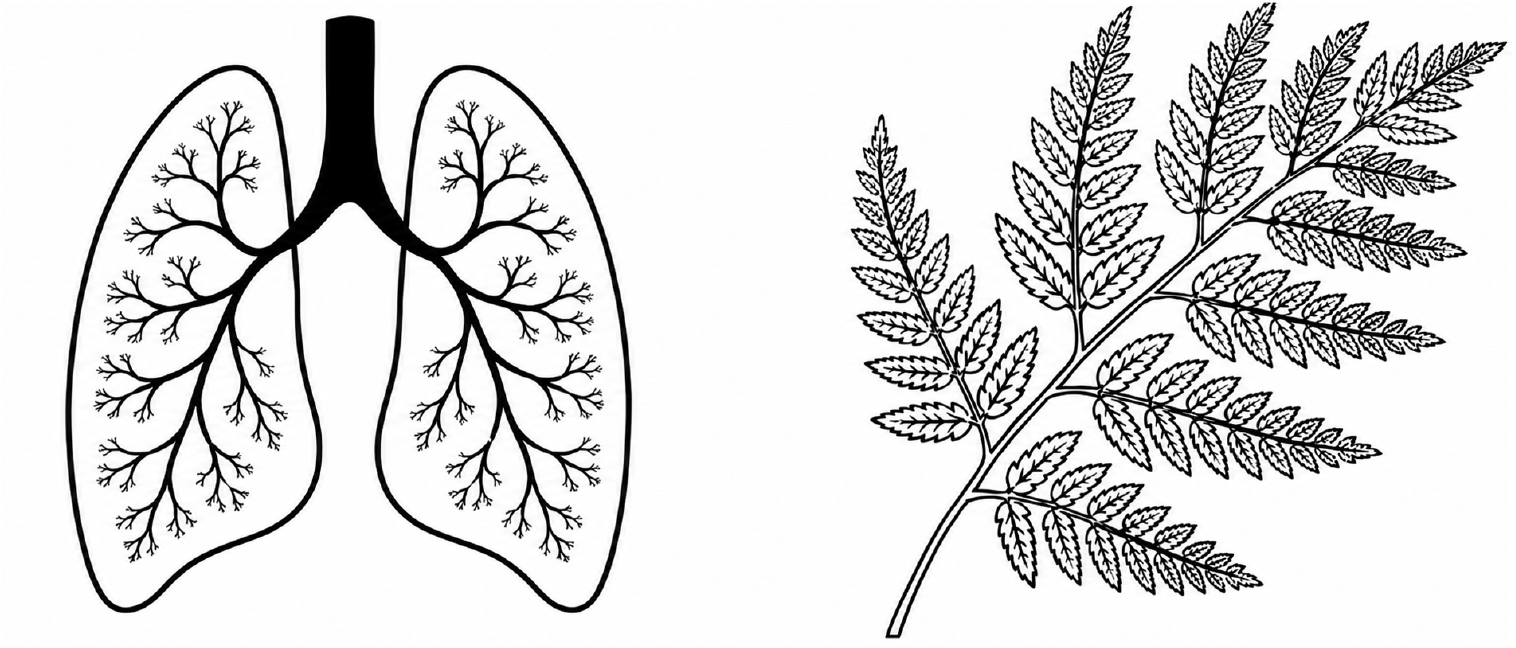
Illustration of self-similarity in nature. The structures shown here, lungs (left) and a fern leaf (right), exhibit recurring branching patterns across scales.

**Figure 4:**
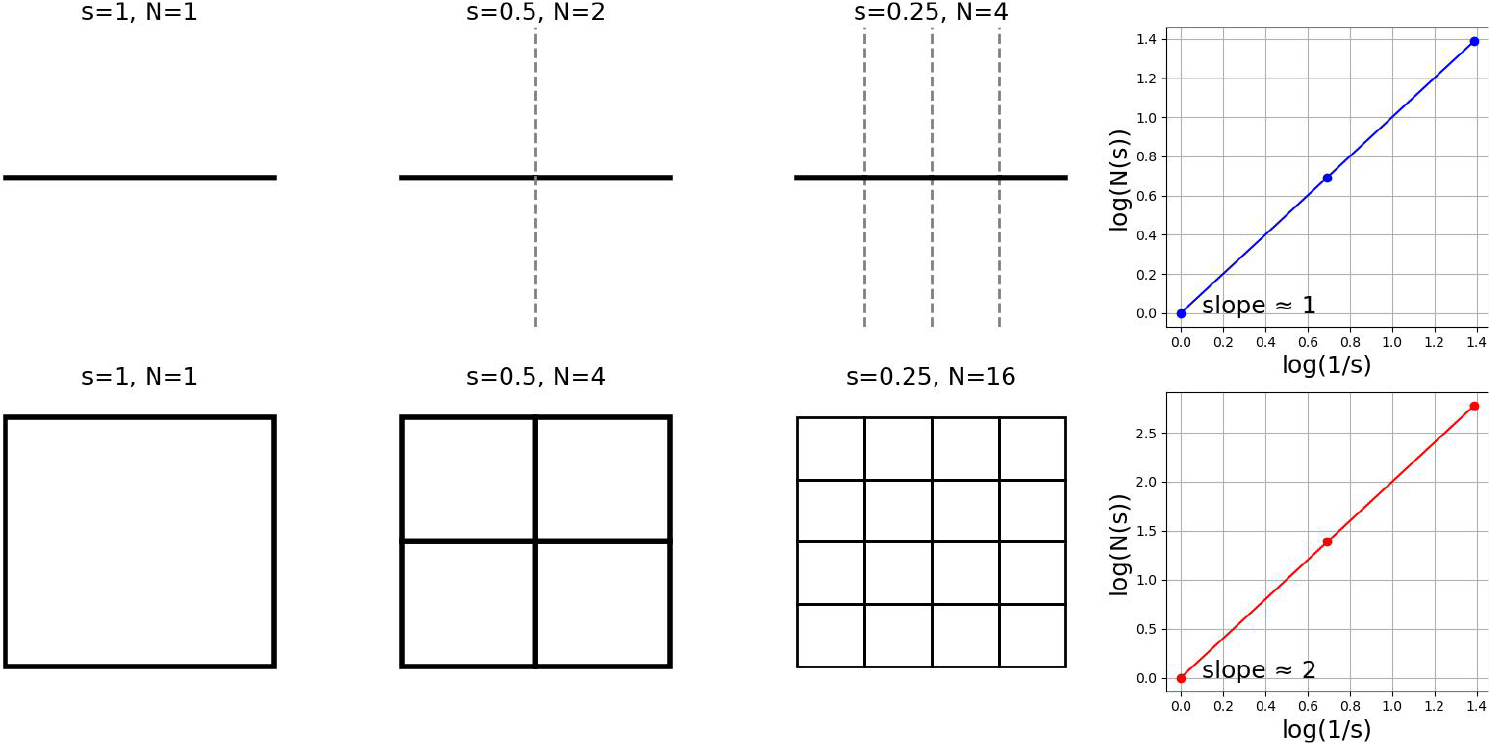
Illustration of the box-counting method. As the box size *s* decreases, the number of boxes *N* (*s*) required to cover the object increases. The log-log plot of log *N* (*s*) versus log(1*/s*) yields a slope corresponding to the fractal dimension. The top row illustrates a one-dimensional line (slope ≈ 1), and the bottom row shows a two-dimensional square (slope ≈ 2).

In classical Euclidean geometry, dimension is integer-valued: a line is one-dimensional, a plane is two-dimensional, and a volume is three-dimensional. Many natural objects, however, are not well described by integer dimensions. Their geometric complexity is often better captured by a fractal dimension, which quantifies how structural detail changes with scale.

A classical way to formalize this idea is through box-counting [15, 16, 17]. One covers the object with boxes of side length *s* and counts the number *N* (*s*) required to cover it. For a one-dimensional object, halving the scale roughly doubles the number of boxes; for a two-dimensional object, it roughly quadruples it. This scaling behavior motivates the box-counting dimension

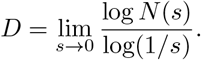

When a linear relationship is observed between log *N* (*s*) and log(1*/s*) over a suitable range of scales, the slope provides an estimate of the dimension. In practice, such scaling relationships are evaluated on a discrete collection of scales, and throughout this work we use logarithmically spaced radius values in order to assess power-law behavior in a balanced manner across multiple orders of magnitude. In the present work, our objective is analogous: we seek to quantify how chromatin interaction networks scale as the observation scale varies, thereby assessing whether their organization is compatible with fractal-like behavior.

Several notions of fractal dimension have been introduced in the literature. The Hausdorff dimension is a classical set-based definition, the Hurst exponent is widely used for stochastic processes and time series, and box-counting is common in geometric applications. In this paper, we focus on two graph-based approaches that are suitable for weighted chromatin interaction networks: the correlation dimension and the sandbox dimension.

### 2.2.1 Fractal Dimensions of Networks: The Correlation Approach

Box-counting procedures on networks are often computationally demanding, especially for large weighted graphs [18]. An alternative is the correlation dimension, which characterizes scaling behavior through the distribution of pairwise graph distances [19].

Let *G* denote a chromatin interaction network. We say that *G* has correlation dimension *d*_*C*_ if the correlation sum *C*(*s*) satisfies

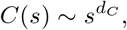

over an appropriate range of scales *s*. The correlation sum is defined by

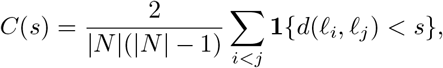

where |*N*| is the number of nodes in the graph, *ℓ*_*i*_ and *ℓ*_*j*_ denote genomic loci, *d*(*ℓ*_*i*_, *ℓ*_*j*_) is the graph distance between them, and **1** {·} is the indicator function. Thus, *C*(*s*) measures the proportion of node pairs whose graph distance is smaller than the threshold *s*.

If the plot of log *C*(*s*) against log *s* is approximately linear over a nontrivial range of scales, then the slope provides an estimate of *d*_*C*_. In this sense, the correlation dimension captures how quickly neighborhoods in the graph fill out as the distance threshold increases, and therefore provides a quantitative summary of network self-similarity.

### 2.2.2 Fractal Dimensions of Networks: The Sandbox Approach

A single fractal dimension may not always capture the full heterogeneity of a complex network. This motivates the use of generalized dimensions *D*_*q*_, which describe scaling behavior through moments of the mass distribution across scales. For a network *G* with a covering *ℬ* (*s*) by boxes of size at most *s*, if *p*_*j*_(*s*) denotes the fraction of total mass contained in box *B*_*j*_, then for *q* ≠ 1 the generalized dimension is defined by

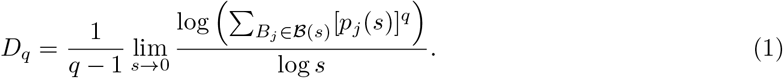

This definition includes several standard notions as special cases. In particular, *q* = 0 corresponds to the box-counting dimension, *q* = 1 leads to the information dimension, and *q* = 2 is related to the correlation dimension.

For large and irregular networks, however, exact minimal coverings are difficult to compute. The sandbox method avoids this difficulty by estimating the scaling relation through local neighborhoods centered at sampled nodes. Let ℒ ⊆ *N* denote a subset of sampled loci. For each *ℓ*_*i*_ ∈ ℒ, define *M* (*ℓ*_*i*_, *s*) as the number of nodes within graph distance at most *s* from *ℓ*_*i*_. The corresponding normalized mass is

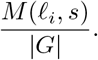

Averaging the (*q* − 1)st power of this quantity over the sampled centers gives

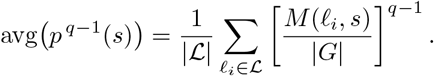

Under the usual scaling assumption, this average behaves as

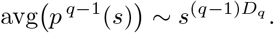

This motivates the sandbox estimator

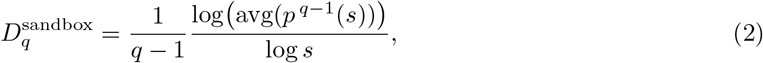

for sufficiently small scales *s*. In practice, the sandbox approach replaces the global covering problem by repeated local measurements of graph mass growth, which makes it attractive for large weighted chromatin networks.

Figure 5 illustrates several example networks with distinct fractal behavior under the correlation and sandbox approaches.

**Figure 5:**
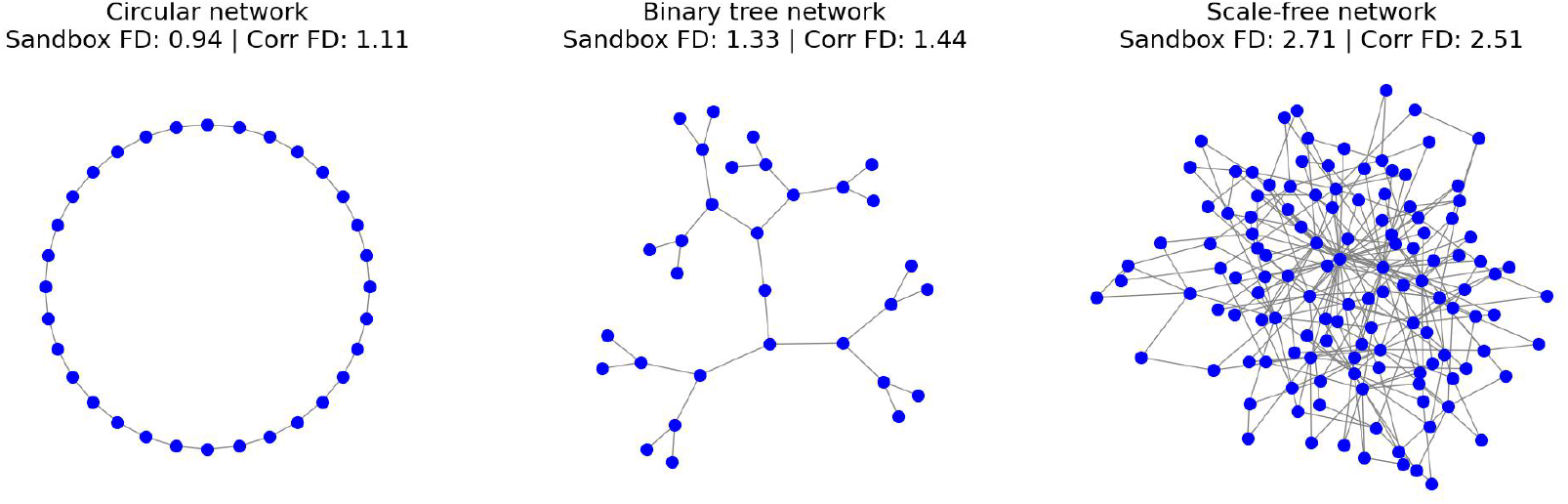
Examples of networks with distinct fractal dimensions as computed by the correlation and sandbox approaches. Left: a circular network. Middle: a binary tree. Right: a scale-free network.

## 3 Validation

Before applying the proposed framework to Hi-C chromatin interaction networks, we assess its behavior on controlled network examples. The objective of this section is threefold. First, we verify that the proposed estimators distinguish networks with clear fractal-like scaling from networks that do not exhibit such behavior. Second, we quantify the uncertainty induced by sampling choices in the correlation and sandbox approaches. Third, we study the computational cost of both methods as the network size increases.

### 3.1 Binary tree vs ring of Erdős–Rényi clusters

We begin with a qualitative and quantitative validation on two synthetic networks with contrasting structural organization. The first network is a balanced binary tree, which provides a canonical example of recursive branching and explicit multiscale hierarchy. The second is a ring of Erdős–Rényi clusters, constructed by connecting several locally random modules in a circular arrangement. This second network exhibits strong local connectivity within clusters together with a simple global cyclic organization, but it does not possess the recursive self-similar structure of the hierarchical tree. The two synthetic networks are displayed in Figure 6.

**Figure 6:**
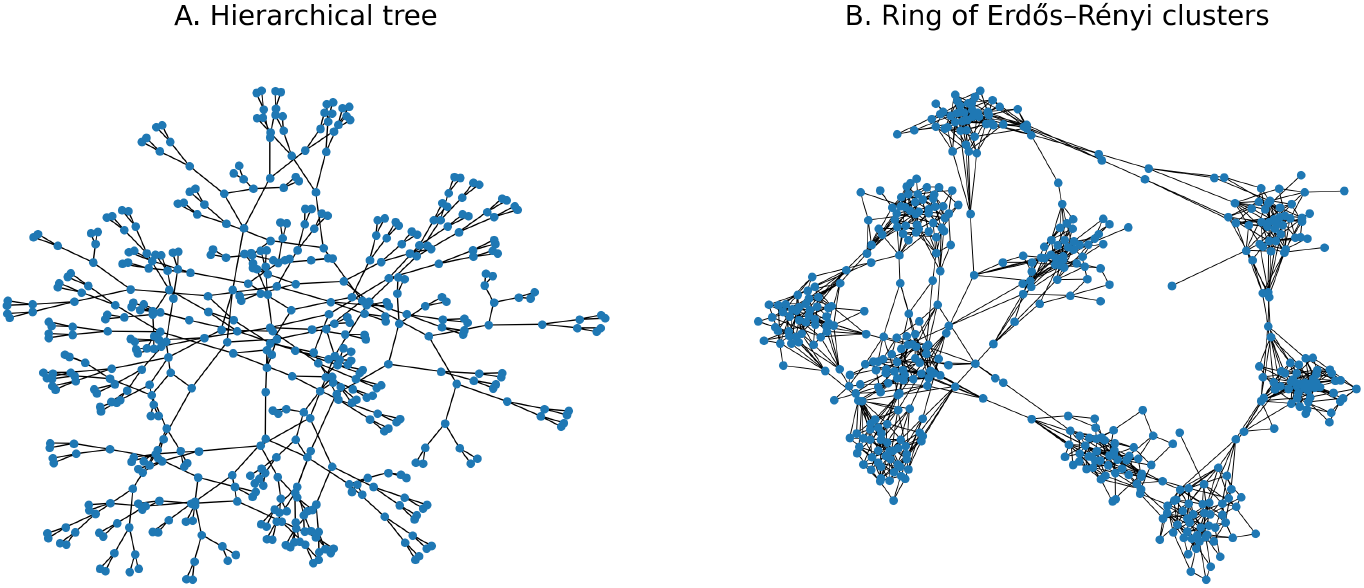
Synthetic networks used for validation. Panel A shows a hierarchical tree with recursive branching structure. Panel B shows a ring of Erdős–Rényi clusters, constructed from locally connected random modules arranged in a cyclic topology.

For each network, we estimate the fractal dimension using both the correlation and sandbox approaches. More precisely, for the correlation method we examine the log-log relationship between the correlation sum *C*(*s*) and the scale parameter *s*, while for the sandbox method we study the log-log scaling of the averaged mass-radius function. If a network exhibits meaningful fractal-like scaling, one expects these curves to display an approximately linear regime over a nontrivial range of scales. In contrast, for a network whose structural organization varies substantially across scales, the scaling curves are not expected to remain close to linear over the full range.

The hierarchical tree provides a clear positive-control example. As shown in Figure 7, both the correlation and sandbox curves are close to linear across a substantial range of scales, indicating stable scaling behavior consistent with self-similar organization. The corresponding estimated dimensions are also highly concordant across methods, with correlation and sandbox estimates both close to 3. This agreement supports the ability of the proposed estimators to recover coherent scaling behavior when the underlying network has an explicitly hierarchical structure.

**Figure 7:**
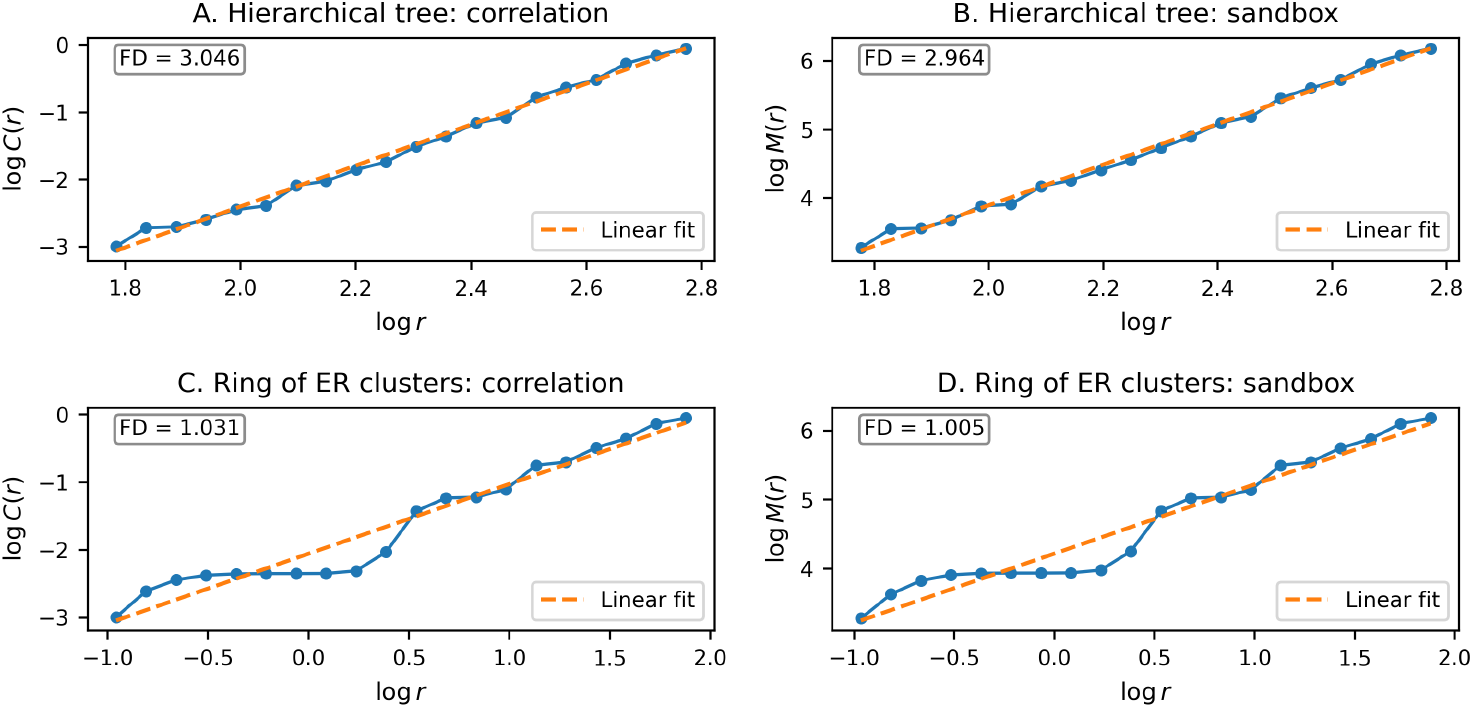
Log-log scaling curves for the synthetic networks in Figure 6 under the correlation and sandbox approaches. Panels A and B correspond to the hierarchical tree, which exhibits an approximately linear scaling regime and consistent dimension estimates across methods. Panels C and D correspond to the ring of Erdős–Rényi clusters, whose scaling curves vary more substantially across scales and indicate weaker evidence of self-similar organization.

**Figure 8:**
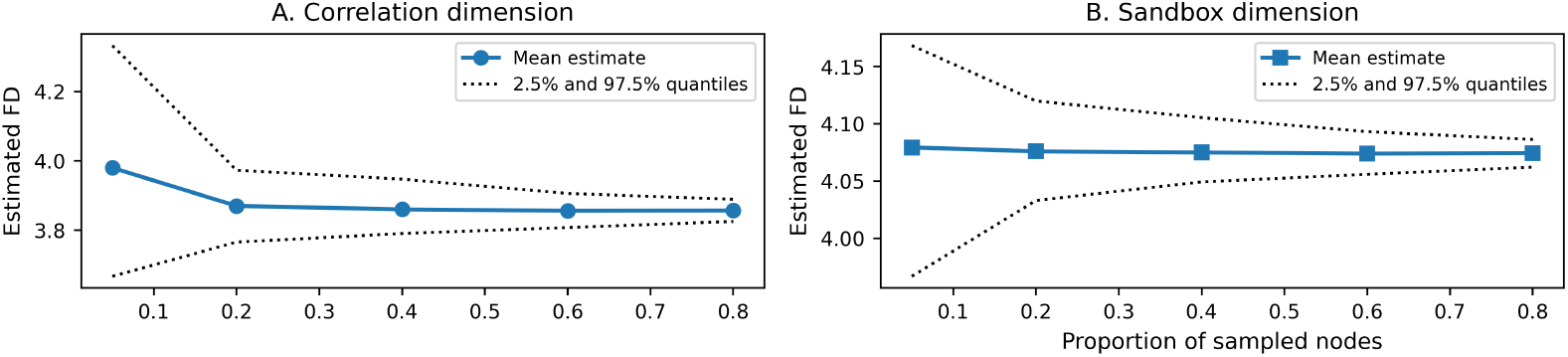
Uncertainty of the estimated fractal dimension as a function of the proportion of sampled nodes, for the correlation and sandbox approaches.

By contrast, the ring of Erdős–Rényi clusters behaves differently. Although this network has pronounced local structure, its global organization is governed by a cyclic arrangement of clusters rather than by recursive branching. Accordingly, the scaling curves in Figure 7 do not exhibit a comparably stable linear regime. In particular, the slopes vary across scales. At small scales, the curves are steeper because the radius expansion is largely confined within individual Erdős–Rényi clusters, which are relatively compact and well connected, leading to rapid neighborhood growth. At larger scales, the expansion increasingly depends on the much sparser inter-cluster connections along the ring, so the growth rate slows substantially and the slope decreases. Over intermediate scales, the curves may transiently approach a more linear behavior, but this regime is neither as stable nor as extensive as in the hierarchical tree. The resulting estimated dimensions are close to 1, reflecting the dominance of the large-scale cyclic organization rather than a genuinely self-similar multiscale structure. The associated scaling curves are shown in Figure 7.

Taken together, these examples show that the correlation and sandbox estimators recover coherent scaling behavior on a network with explicit hierarchical organization, while producing markedly different scaling diagnostics on a network with strong local clustering but without recursive multiscale structure.

### 3.2 Uncertainty in FD estimates

Both the correlation and sandbox approaches may depend on the number of sampled nodes used during estimation. We therefore assess the stability of the estimated fractal dimension as a function of the sampling proportion. For a fixed network size and a fixed network structure, we consider a collection of sampling proportions

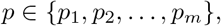

where *p* denotes the proportion of nodes selected for estimation. For each value of *p*, we repeat the estimation procedure multiple times and summarize the resulting distribution of fractal-dimension estimates. This allows us to evaluate how the estimator variability decreases as the number of sampled nodes increases.

The main quantity of interest is the uncertainty width, defined for example by the empirical standard deviation or by the width of an empirical quantile interval across repeated runs. We then plot this uncertainty measure as a function of the sampling proportion for both methods.

Overall, this experiment provides practical guidance for selecting the sampling parameter in large networks and supports the use of node subsampling as a computationally efficient approximation strategy when full estimation is costly. More precisely, for the correlation estimator, sampling 60% of the nodes yields a relative empirical error of approximately 2.54%, while sampling 80% reduces it to approximately 1.65%. The sandbox estimator is even more stable, with relative empirical error approximately 0.91% at 60% sampling and approximately 0.59% at 80%. These results indicate that, for sufficiently large networks, accurate fractal-dimension estimates can often be obtained from a substantial subsample of nodes, thereby reducing computational cost while preserving high estimation accuracy.

### 3.3 Computational complexity of fractal dimension estimation

Both the correlation and sandbox estimators rely on repeated shortest-path computations in weighted graphs. In our implementation, graph distances are obtained through repeated calls to Dijkstra’s single-source shortest-path algorithm. More precisely, for each sampled node, we compute its shortest-path distance to all reachable nodes in the graph using the edge weights defined by the network construction. This distance computation is the main computational bottleneck in both approaches [19, 18].

For a weighted graph with *N* nodes and *E* edges, a single run of Dijkstra’s algorithm has complexity on the order of *O*((*N* + *E*) log *N*), or *O*(*E* log *N*) in sparse graphs. If *k* sampled nodes are used in the estimation procedure, then both the correlation and sandbox methods require approximately *k* such shortest-path computations. Consequently, the dominant computational cost is of order

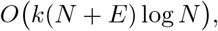

up to additional lower-order operations associated with computing the correlation sums or sandbox masses once the distances have been obtained. When the sampling proportion is *p*, one typically has *k* ≈ *pN*, so the computational burden increases with both the graph size and the proportion of nodes used in estimation.

To study this effect empirically, we generated connected Erdős–Rényi graphs with weighted edges and measured the runtime of the correlation and sandbox estimators as the number of nodes increased. Specifically, we considered graph sizes

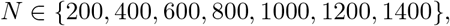

used edge probability 0.1, assigned random edge weights uniformly on [0.1, 1.0], and evaluated both estimators with 20 logarithmically spaced radii. We examined two sampling proportions, *p* = 0.2 and *p* = 0.6, in order to quantify the computational effect of using a larger subset of nodes. The resulting runtimes are displayed in Figure 9.

**Figure 9:**
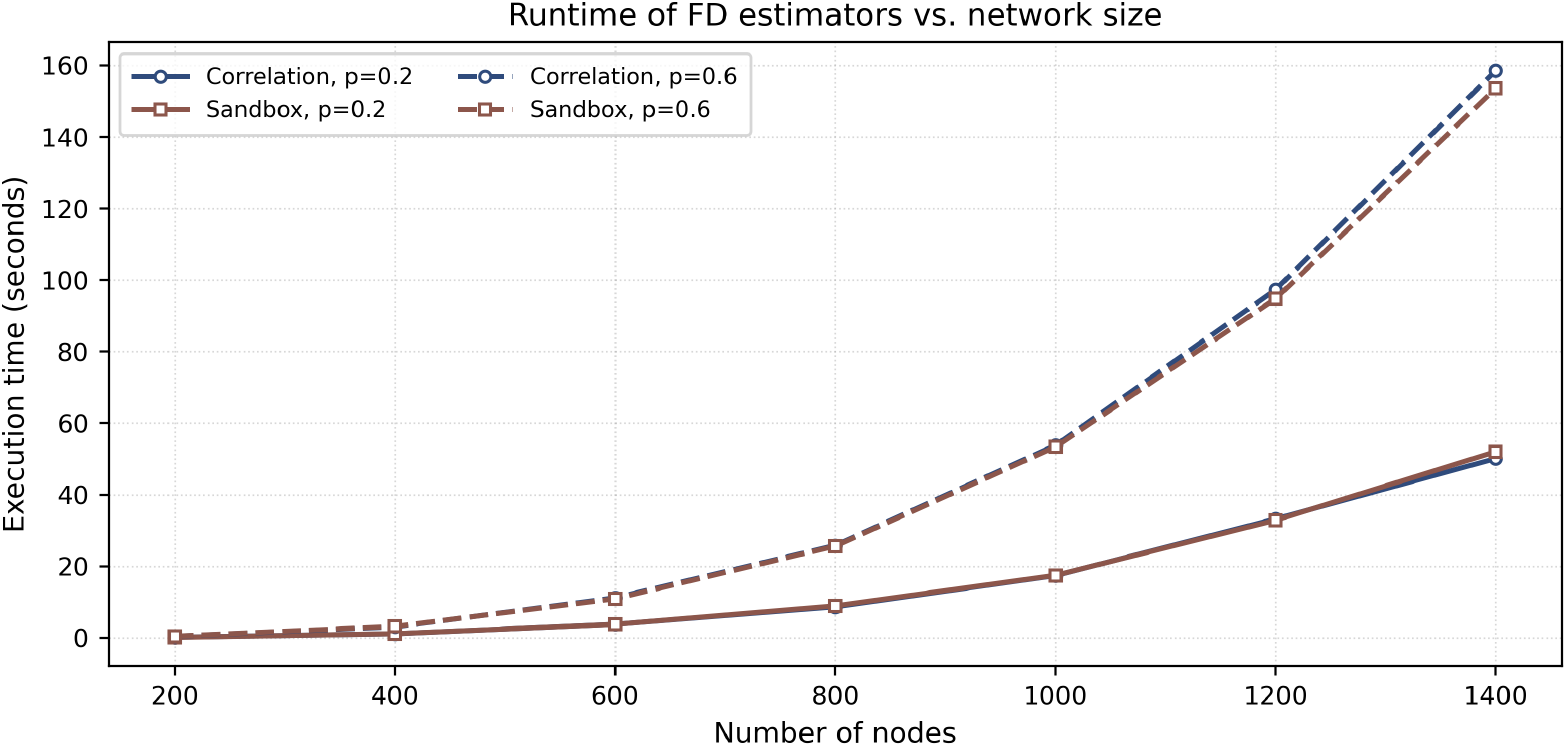
Empirical runtime of the correlation and sandbox fractal-dimension estimators as a function of network size for two node sampling proportions. Colors distinguish the two estimation methods, while line styles indicate the sampling proportion. Larger sampling proportions lead to substantially increased computational cost for both estimators.

As expected, the runtime increases rapidly with network size for both estimators. Moreover, the computational burden is substantially higher when the sampling proportion increases from 0.2 to 0.6, reflecting the larger number of shortest-path computations required. In our experiments, the correlation and sandbox methods exhibited very similar runtimes, which is consistent with the fact that both are dominated by repeated calls to Dijkstra’s algorithm. Thus, from a computational perspective, the principal factor controlling runtime is not so much the choice between the two estimators, but rather the graph size and the number of sampled nodes.

This issue is directly relevant for Hi-C analysis. For the largest human chromosomes, whose lengths are on the order of 250 Mbp, the corresponding contact matrices already become substantial at moderate resolution: at 25 kb resolution, one obtains on the order of 10^4^ bins, which is still manageable on a standard laptop but already leads to long processing times for distance-based fractal-dimension estimation. At 5 kb resolution, the same chromosome would be represented by on the order of 5 *×* 10^4^ bins, making full chromosome-wide computation far more demanding in both memory and runtime. Thus, obtaining reliable estimates requires balancing the proportion of sampled nodes against the computational cost: larger sampling proportions generally improve stability, but they also substantially increase the number of shortest-path computations.

## 4 Application

The proposed framework is applied to intrachromosomal Hi-C data from the IMR90 human cell line. Since these data consist of contact matrices that record interaction frequencies between genomic loci, they provide a natural multiscale representation of chromatin organization and are therefore well suited for fractal analysis. Our objective is to assess whether the resulting chromatin contact networks exhibit scaling behavior consistent with fractal organization, and to study how the corresponding fractal-dimension estimates vary across chromosomes and genomic scales.

To illustrate the type of data under consideration, Figure 10 displays representative Hi-C contact maps for chromosomes 1–4 at two resolutions, 500 kb and 100 kb. For visualization purposes, the matrices are shown after the transformation log(count + 1), which improves contrast and makes both strong and moderate interaction patterns visible. As expected, the maps exhibit a pronounced diagonal structure, reflecting the fact that nearby loci interact more frequently, together with block and band patterns that suggest nontrivial multiscale chromatin organization. These visual features motivate the use of fractal descriptors. In particular, the apparent repetition of structural motifs across scales suggests approximate scaling behavior, which we investigate quantitatively through the proposed estimators.

**Figure 10:**
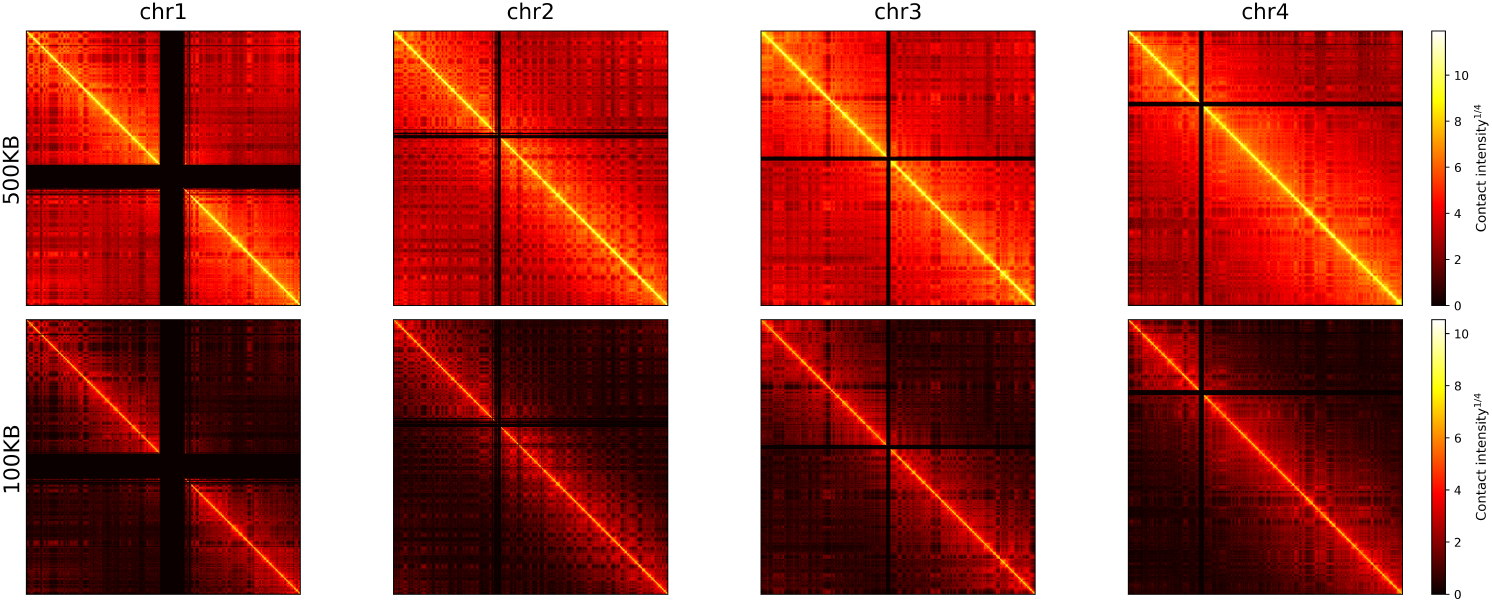
Illustrative Hi-C contact maps for chromosomes 1–4 at 500 kb and 100 kb resolution in the IMR90 cell line. For visualization, contact counts are displayed on the log(count + 1) scale. The maps reveal clear multiscale structure and strong diagonal organization, consistent with spatial proximity between nearby genomic loci.

### 4.1 Chromosome-level scaling curves and fractal dimensions

The fractal dimension of each chromosome is estimated using the correlation and sandbox methods described in Section 2. We focus on the 50 kb and 25 kb resolutions, which provide a useful balance between biological detail and computational feasibility. For each chromosome and each method, the estimated scaling curves exhibit a pronounced approximately linear regime on the log–log scale, supporting the presence of fractal-like behavior in chromatin contact organization.

Figure 11 shows the chromosome-wise scaling curves at 50 kb resolution, while Figure 12 shows the corresponding results at 25 kb resolution. In each figure, chromosomes are grouped by estimated FD quartiles. The upper panels correspond to the correlation method and the lower panels to the sandbox method. Across both resolutions, the first group contains chromosomes with the largest estimated dimensions, typically between 2 and 3, indicating highly compact structures that occupy a substantial portion of three-dimensional space. In contrast, the lowest-ranked group contains chromosomes with dimensions closer to 1, suggesting more open or less space-filling conformations.

**Figure 11:**
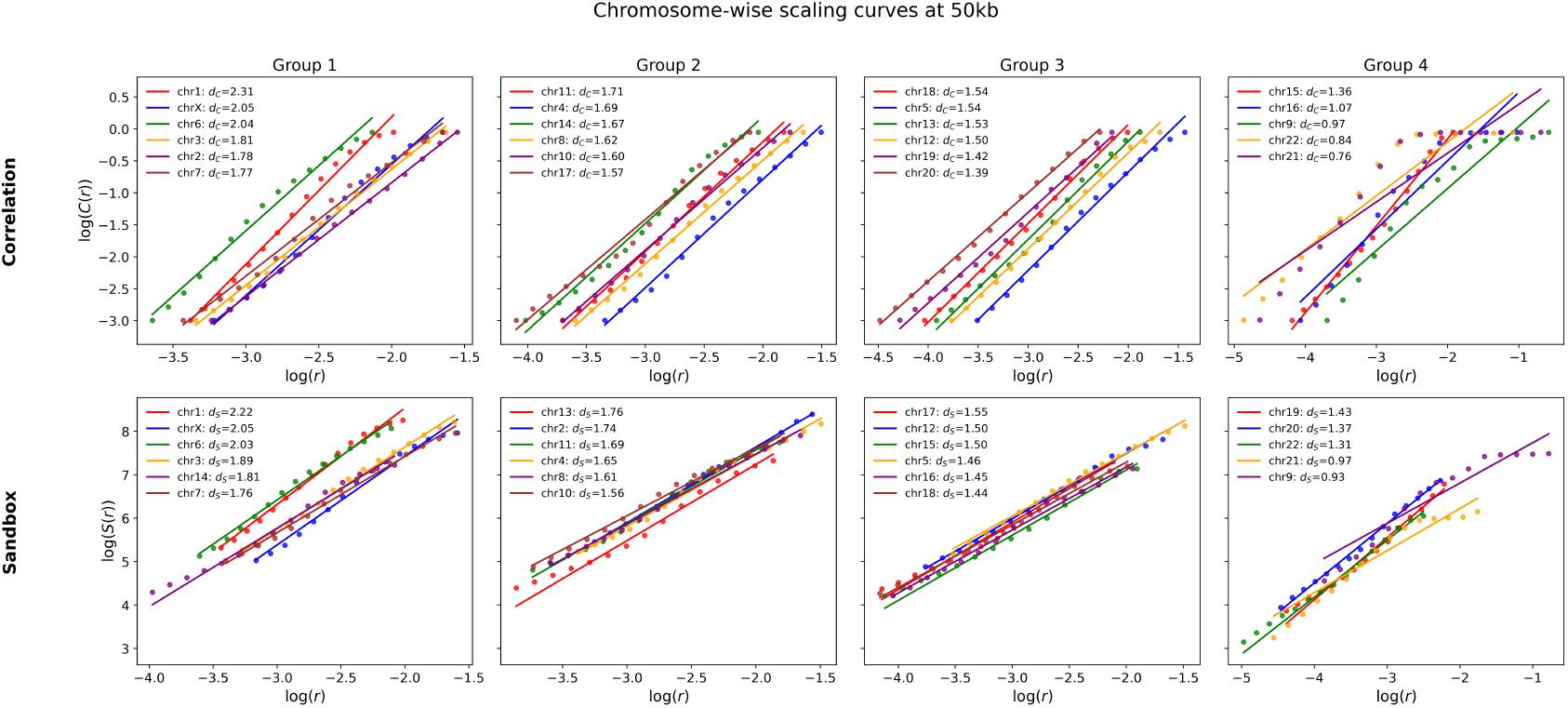
Chromosome-wise scaling curves at 50 kb resolution. The top row corresponds to the correlation method and the bottom row to the sandbox method. Chromosomes are grouped according to estimated fractal-dimension quartiles. The approximately linear log–log scaling supports fractal-like organization of chromatin contact maps.

**Figure 12:**
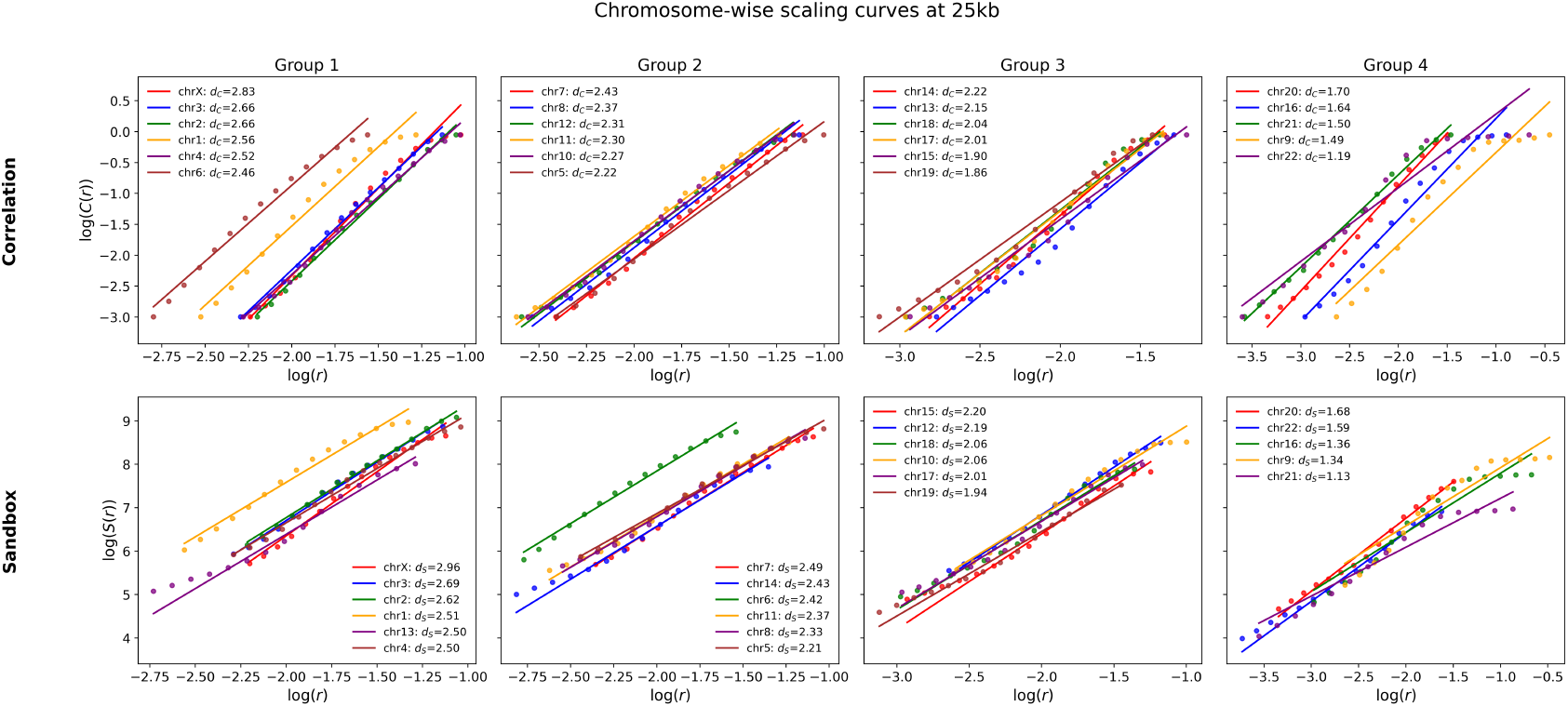
Chromosome-wise scaling curves at 25 kb resolution. The same qualitative pattern as at 50 kb is observed, with clear approximate linear scaling across chromosomes and substantial variability in estimated fractal dimension.

A consistent pattern emerges across resolutions: chromosomes 1, 2, and X tend to have the largest estimated fractal dimensions, whereas chromosomes 21 and 22 tend to have the smallest. This ranking is biologically plausible, as the largest chromosomes display more complex and spatially extensive folding than the shortest chromosomes.

### 4.2 Relationship between chromosome size and fractal dimension

To quantify the association between chromosome size and geometric complexity, we regress chromosome-level FD estimates against chromosome length measured in number of bins at the corresponding resolution. Figure 13 presents the results at 50 kb resolution, and Figure 14 presents the results at 25 kb resolution, with the correlation and sandbox estimates shown side by side.

**Figure 13:**
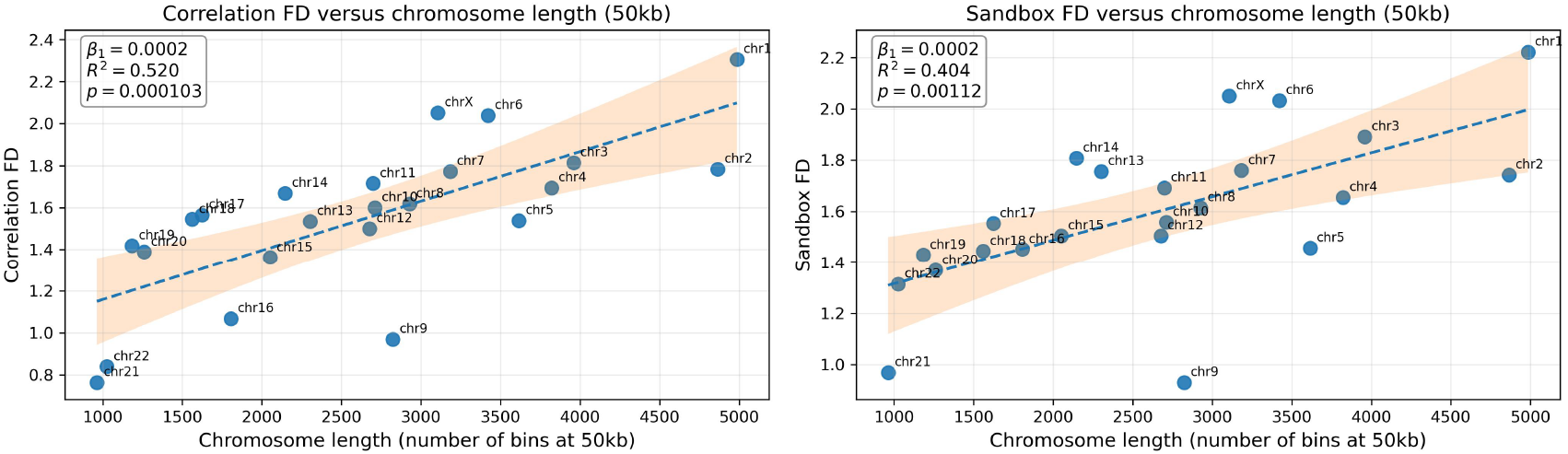
Relationship between chromosome length and estimated fractal dimension at 50 kb resolution. Left: correlation FD. Right: sandbox FD. In both cases, longer chromosomes tend to exhibit higher estimated dimensions.

**Figure 14:**
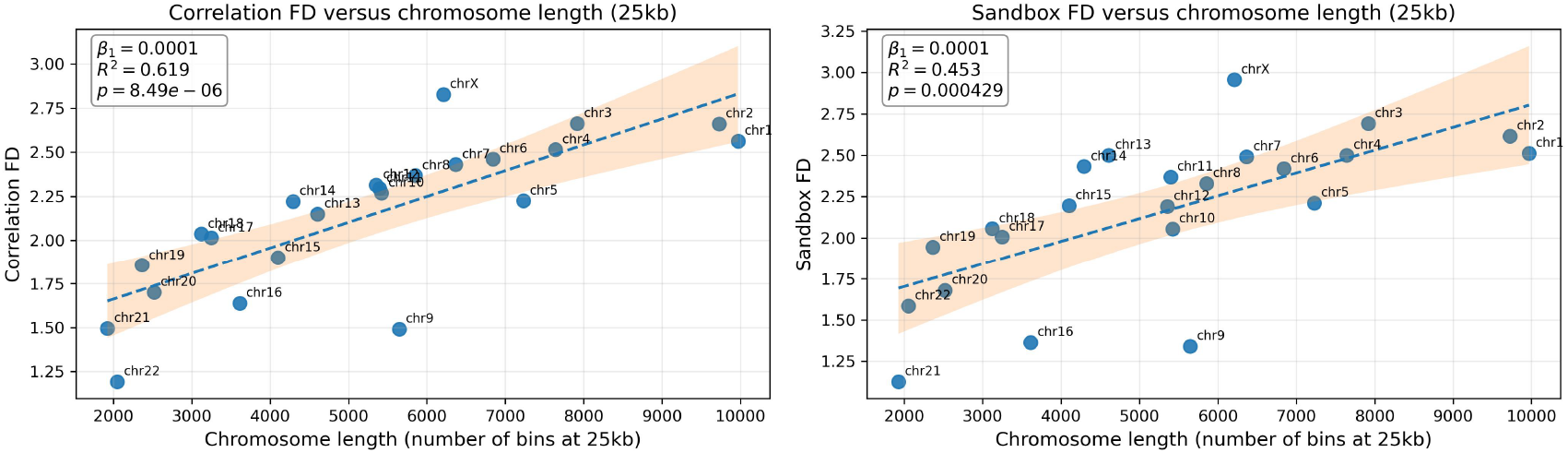
Relationship between chromosome length and estimated fractal dimension at 25 kb resolution. The positive association is stronger than at 50 kb, especially for the correlation-based estimator.

A clear positive relationship is observed: larger chromosomes tend to have higher estimated fractal dimensions, with values approaching 3 for the largest chromosomes and closer to 1 for the shortest chromosomes. This suggests that chromosome size is strongly associated with the degree of space-filling chromatin folding.

At 50 kb resolution, the sandbox estimates yield *R*^2^ ≈ 0.40 with *p* ≈ 10^−3^, while the correlation estimates yield *R*^2^ ≈ 0.52 with *p* ≈ 10^−4^. At 25 kb resolution, the association becomes even stronger: the sandbox estimates give *R*^2^ ≈ 0.45 with *p* ≈ 4 *×* 10^−4^, and the correlation estimates give *R*^2^ ≈ 0.62 with *p <* 10^−7^. Overall, the correlation method appears to produce the strongest size–FD relationship, though both estimators lead to the same qualitative conclusion.

### 4.3 Sliding-window analysis at 5 kb resolution

A single global FD per chromosome is informative, but it may conceal important local heterogeneity. To investigate spatial variation in chromatin organization along the chromosome, we perform a sliding-window analysis at 5 kb resolution. For each chromosome, we extract overlapping diagonal submatrices and estimate local fractal dimension using both the correlation and sandbox methods. This provides a profile of local structural complexity along genomic position.

Figures 15, 16, and 17 show representative examples for chromosomes 1, 9, and 10, respectively. Several important features are visible. First, the estimated local FD is clearly not homogeneous along the chromosome; instead, it evolves substantially from one region to another, indicating marked spatial variation in chromatin folding complexity. Second, some genomic regions display missing or unstable values, especially around centromeric regions where sequencing and alignment are notoriously difficult. This is particularly pronounced for chromosome 9, which exhibits a large region with limited information, consistent with its prominent centromeric structure. Third, chromosome 10 provides an example where some regions display markedly higher FD than others, suggesting alternating domains of more compact versus more open chromatin organization.

**Figure 15:**
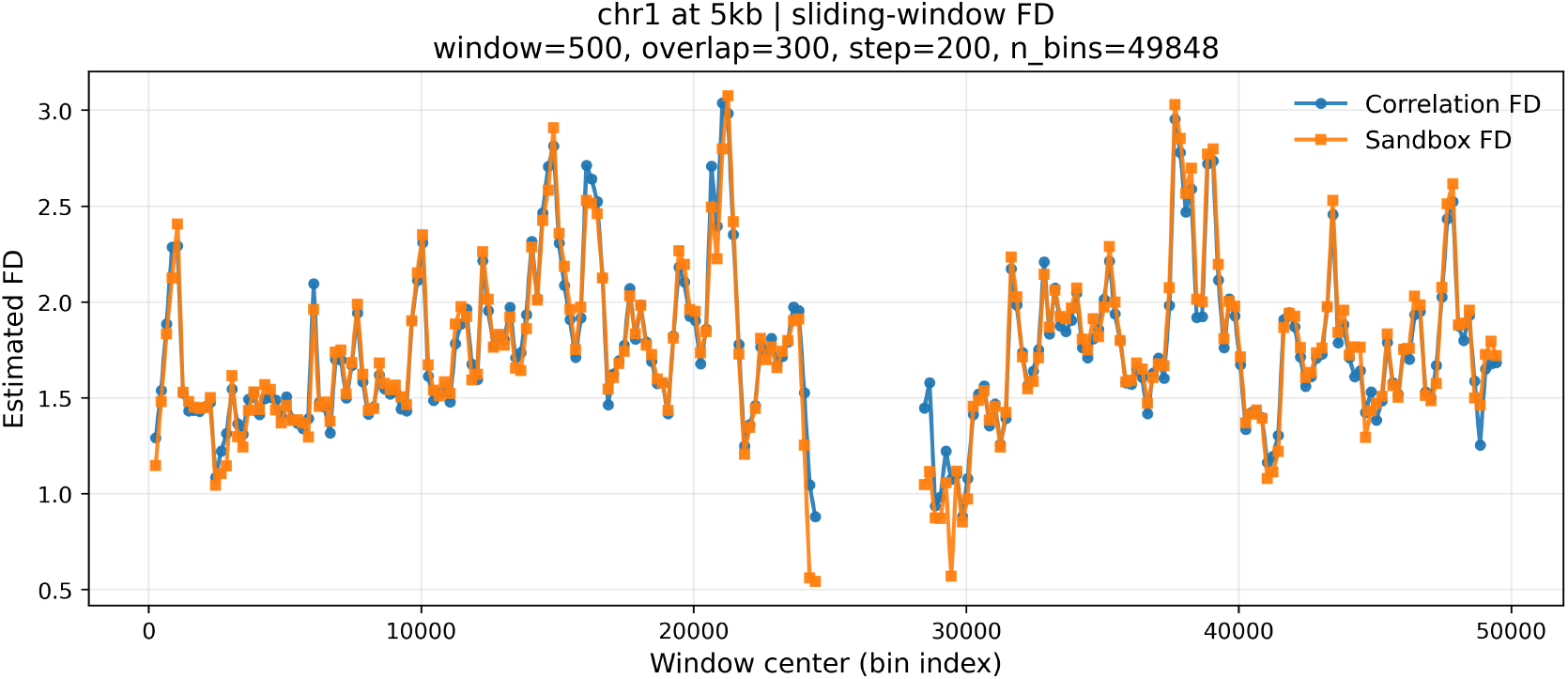
Sliding-window fractal-dimension analysis for chromosome 1 at 5 kb resolution, using window size 500 bins and overlap 300 bins. The local FD varies substantially along the chromosome, showing that chromatin organization is not homogeneous even within a single chromosome.

**Figure 16:**
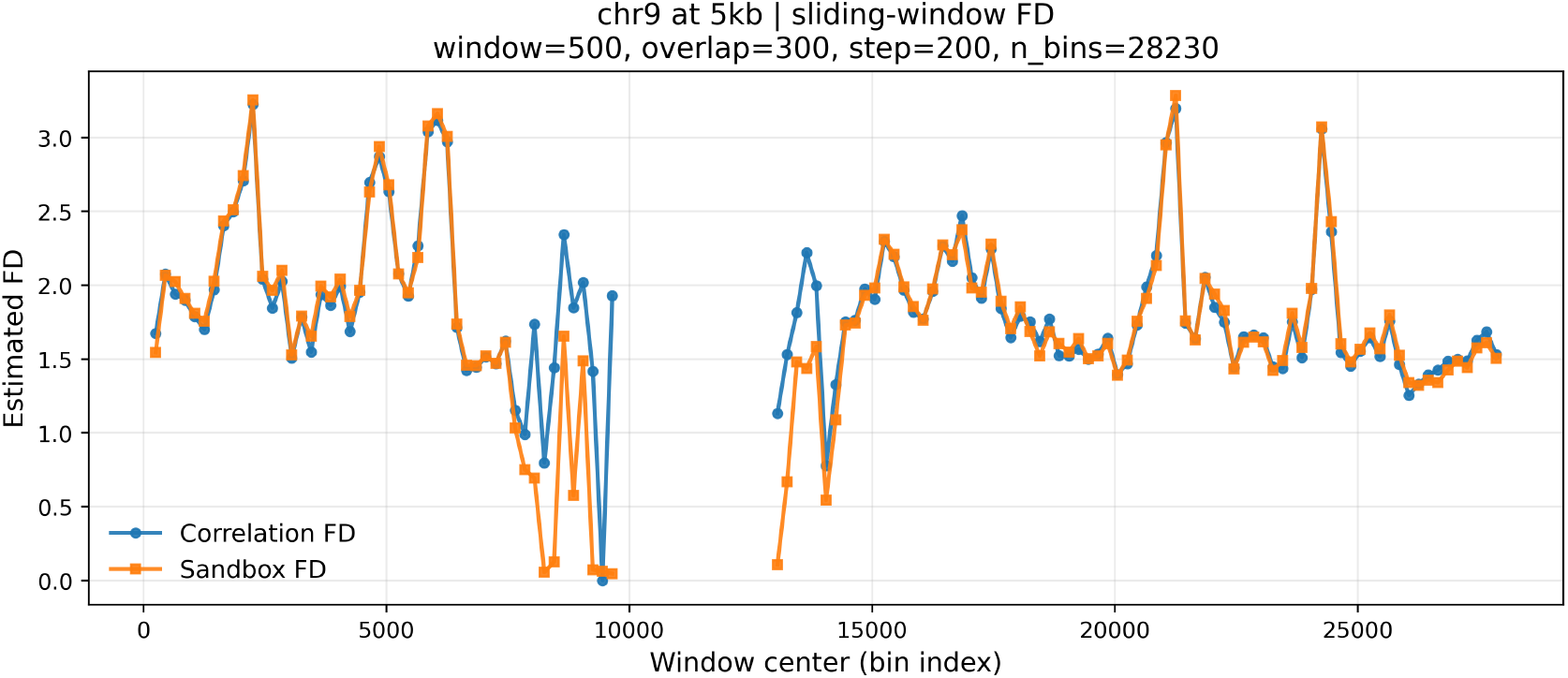
Sliding-window fractal-dimension analysis for chromosome 9 at 5 kb resolution. A broad region of missing or unstable estimates is visible, consistent with the presence of a large centromeric region and limited mappability.

**Figure 17:**
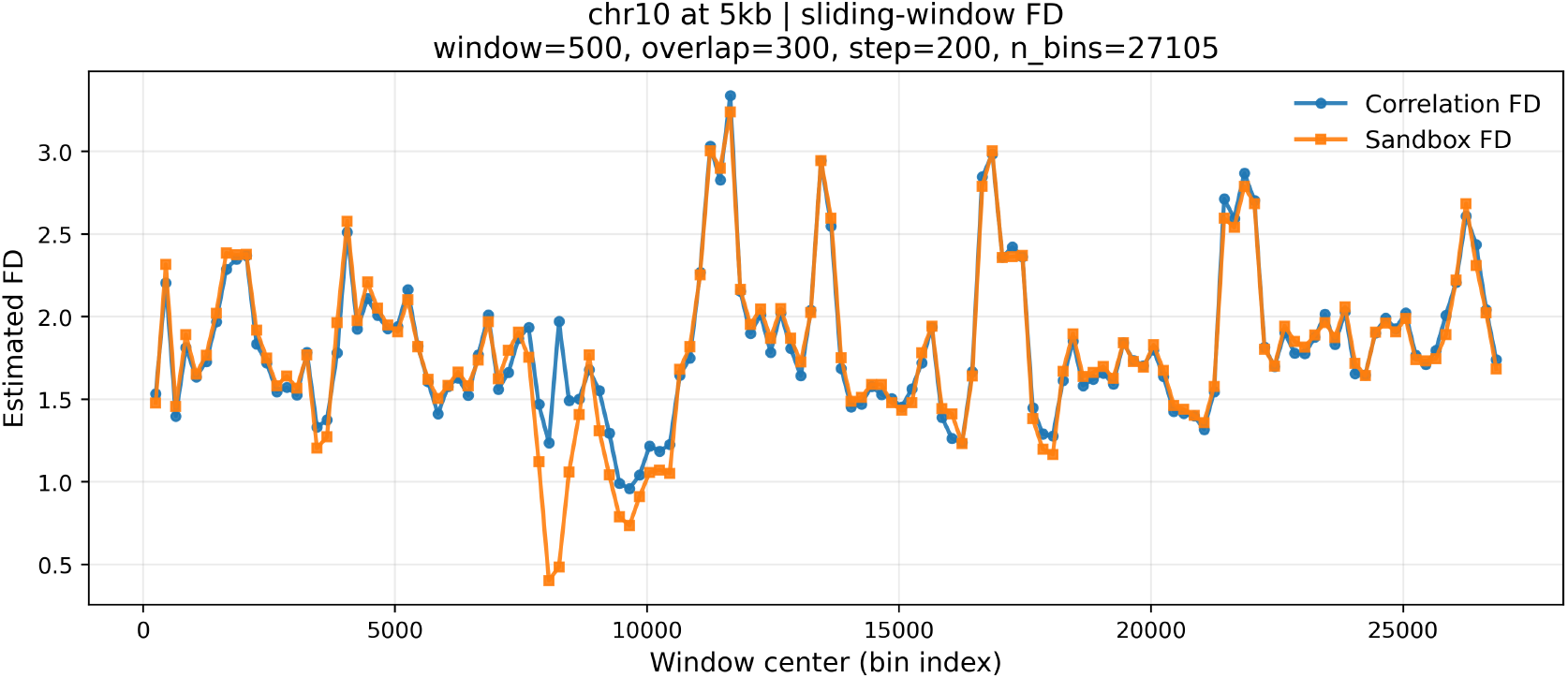
Sliding-window fractal-dimension analysis for chromosome 10 at 5 kb resolution. Distinct regions of higher and lower local FD are evident, suggesting alternating domains of relatively compact and relatively open chromatin organization.

These observations show that local fractal analysis complements the global chromosome-level summary: while overall FD captures broad structural differences between chromosomes, the sliding-window approach reveals substantial within-chromosome heterogeneity that would otherwise be obscured.

Overall, the application demonstrates that Hi-C contact maps exhibit robust scaling behavior compatible with fractal organization, that chromosome-level fractal dimension is strongly associated with chromosome size, and that local FD analysis reveals substantial heterogeneity along individual chromosomes. Together, these results support the usefulness of fractal-dimension estimators as interpretable geometric summaries of chromatin conformation.

## 5 Conclusion

In this paper, we introduced a network-based framework to study the fractal organization of chromatin from Hi-C data. By representing chromosomes as weighted interaction networks and estimating their fractal dimension using both the correlation and sandbox approaches, we quantified self-similar scaling in chromatin architecture across genomic scales. Our results show that fractal dimensions vary substantially across chromosomes: longer chromosomes tend to have higher dimensions, often approaching 3, consistent with a more compact and space-filling three-dimensional organization, whereas shorter chromosomes tend to have lower dimensions, often between 1 and 1.5, suggesting a simpler and more open structure. The sliding-window analysis further revealed that fractal organization is not homogeneous within chromosomes, but evolves across genomic position. Overall, these findings highlight the relevance of fractal analysis as an interpretable and informative tool for studying multiscale genome organization.

A practical limitation of the present framework is that full chromosome-wide fractal-dimension estimation becomes increasingly challenging at very high resolution. As the bin size decreases, the number of loci grows rapidly, which leads to much larger contact matrices and substantially higher computational cost for reliable estimation. Developing more scalable algorithms, sparse implementations, or approximation strategies for high-resolution Hi-C networks therefore remains an important methodological direction. Beyond the present application, the proposed framework opens several promising directions for future research. Fractal-dimension estimates could be compared across different genomic regions, cell types, tissues, developmental stages, or disease conditions in order to assess how chromatin compaction and folding complexity vary across biological contexts. In particular, local fractal-dimension profiles may help identify regions with distinct structural organization and may provide a useful complement to more classical descriptors of chromatin architecture.

One particularly interesting possibility concerns deviations from the global relationship between chromosome length and fractal dimension. Such deviations may reflect biologically meaningful departures from the typical scaling pattern, potentially related to structural constraints such as chromatin knotting. As a polymer, chromatin is in principle susceptible to knotting, and the tendency to form knots is influenced by its length, stiffness, and effective thickness. At the same time, proteins such as histones, condensins, and CTCF impose structural constraints that can limit or reshape this tendency. Understanding whether departures from the observed FD–length trend are associated with such topological constraints could provide an interesting bridge between fractal scaling, chromatin compaction, and the topology of genome organization.

## References

[1] T. Misteli, “The self-organizing genome: Principles of genome architecture and function,” Cell, vol. 183, no. 1, pp. 28–45, 2020.

[2] A. Travers and G. Muskhelishvili, “DNA structure and function,” The FEBS Journal, vol. 282, 2015.

[3] G. Hudson, A. Gomez-Duran, I. J. Wilson, and P. F. Chinnery, “Recent mitochondrial dna mutations increase the risk of developing common late-onset human diseases,” PLoS Genet, vol. 10, p. e1004369, May 2014.

[4] S. Samir, “Human DNA mutations and their impact on genetic disorders,” Recent Pat Biotechnol, vol. 18, no. 4, pp. 288–315, 2024.

[5] O. Zaytseva and L. M. Quinn, “DNA conformation regulates gene expression: the MYC promoter and beyond,” BioEssays, vol. 40, 2018.

[6] K. Mondal and S. Chaudhury, “Effect of DNA conformation on the protein search for targets on DNA: A theoretical perspective,” The Journal of Physical Chemistry B, vol. 124, no. 17, pp. 3518–3526, 2020.

[7] B. van Steensel, “Chromatin: constructing the big picture,” EMBO Journal, vol. 30, pp. 1885–1895, May 2011.

[8] V. Bedin, R. L. Adam, B. C. de Sá, G. Landman, and K. Metze, “Fractal dimension of chromatin is an independent prognostic factor for survival in melanoma,” BMC Cancer, vol. 10, p. 260, Jun 2010.

[9] L. M. Almassalha, A. Tiwari, P. T. Ruhoff, Y. Stypula-Cyrus, L. Cherkezyan, H. Matsuda, M. A. Dela Cruz, J. E. Chandler, C. White, C. Maneval, H. Subramanian, I. Szleifer, H. K. Roy, and V. Backman, “The global relationship between chromatin physical topology, fractal structure, and gene expression,” Scientific Reports, vol. 7, no. 1, p. 41061, 2017.

[10] E. R. Mardis, “The impact of next-generation sequencing technology on genetics,” Trends in Genetics, vol. 24, no. 3, pp. 133–141, 2008.

[11] H. Satam, K. Joshi, U. Mangrolia, S. Waghoo, G. Zaidi, S. Rawool, R. P. Thakare, S. Banday, A. K. Mishra, G. Das, and S. K. Malonia, “Next-generation sequencing technology: Current trends and advancements,” Biology, vol. 12, no. 7, 2023.

[12] J.-M. Belton, R. P. McCord, J. H. Gibcus, N. Naumova, Y. Zhan, and J. Dekker, “Hi–c: A comprehensive technique to capture the conformation of genomes,” Methods, vol. 58, no. 3, pp. 268–276, 2012.

[13] K. Emmett, B. Schweinhart, and R. Rabadan, “Multiscale topology of chromatin folding,” in Proceedings of the 9th EAI International Conference on Bio-Inspired Information and Communications Technologies (Formerly BIONETICS), p. 177–180, ICST (Institute for Computer Sciences, Social-Informatics and Telecommunications Engineering), 2016.

[14] S. S. P. Rao, M. H. Huntley, N. C. Durand, E. K. Stamenova, I. D. Bochkov, J. T. Robinson, A. L. Sanborn, I. Machol, A. D. Omer, E. S. Lander, and E. Lieberman Aiden, “A 3d map of the human genome at kilobase resolution reveals principles of chromatin looping,” Cell, vol. 159, no. 7, pp. 1665–1680, 2014.

[15] B. B. Mandelbrot, “How long is the coast of britain? statistical self-similarity and fractional dimension,” Science, vol. 156, pp. 636–638, 1967.

[16] B. B. Mandelbrot, Les objets fractals: Forme, hasard et dimension. Flammarion, 1975.

[17] B. B. Mandelbrot, The Fractal Geometry of Nature. W.H. Freeman and Company, 1982.

[18] P. T. Kovács, M. Nagy, and R. Molontay, “Comparative analysis of box-covering algorithms for fractal networks,” Applied Network Science, vol. 6, 2021.

[19] E. Rosenberg, A Survey of Fractal Dimensions of Networks. Springer, 2018.

